# Mineral density differences between femoral cortical bone and trabecular bone are not explained by turnover rate alone

**DOI:** 10.1101/2020.06.08.141036

**Authors:** Chloé Lerebours, Richard Weinkamer, Andreas Roschger, Pascal R. Buenzli

## Abstract

Bone mineral density distributions (BMDDs) are a measurable property of bone tissues that depends strongly on bone remodelling and mineralisation processes. These processes can vary significantly in health and disease and across skeletal sites, so there is high interest in analysing these processes from experimental BMDDs. Here, we propose a rigorous hypothesis-testing approach based on a mathematical model of mineral heterogeneity in bone due to remodelling and mineralisation, to help explain differences observed between the BMDD of human femoral cortical bone and the BMDD of human trabecular bone. Recent BMDD measurements show that femoral cortical bone possesses a higher bone mineral density, but a similar mineral heterogeneity around the mean compared to trabecular bone. By combining this data with the mathematical model, we are able to test whether this difference in BMDD can be explained by (i) differences in turnover rate; (ii) differences in osteoclast resorption behaviour; and (iii) differences in mineralisation kinetics between the two bone types. We find that accounting only for differences in turnover rate is inconsistent with the fact that both BMDDs have a similar spread around the mean, and that accounting for differences in osteoclast resorption behaviour leads to biologically inconsistent bone remodelling patterns. We conclude that the kinetics of mineral accumulation in bone matrix must therefore be different in femoral cortical bone and trabecular bone. Although both cortical and trabecular bone are made up of lamellar bone, the different mineralisation kinetics in the two types of bone point towards more profound structural differences than usually assumed.

## 1 Introduction

The mineral heterogeneity of bone tissues is the result of remodelling and mineralisation processes, which are known to vary significantly in health and disease and across skeletal sites [7, 27, 36, 61, 47]. Bone remodelling periodically replaces old bone with an unmineralised collagen-rich matrix, which is subsequently infiltrated with a mineral phase. The growth of mineral crystals in this phase gradually confers to bone its stiffness and strength [52, 20]. Regions of bone formed at different times achieve various degrees of mineralisation and are visualised as different grey-level intensities in bone scans obtained by X-ray absorption or quantitative backscattered electron imaging (qBEI) [57, 7, 16, 51, 54, 64, 21]. The heterogeneity of mineral density in a bone scan gives an indication of bone’s renewal history and provides an indirect measure of remodelling and mineralisation processes [10]. Bone mineral density distributions (BMDDs) quantify this mineral heterogeneity experimentally as frequency distributions of calcium content constructed from qBEI scans [58, 61].

In adult trabecular bone, the BMDD is independent of age, ethnicity, sex, and skeletal site (transiliac bone, vertebrae, femoral neck, femoral head, and patella) [58, 61], and the relatively low inter-individual variation has enabled the definition of a reference trabecular BMDD of healthy adults [58]. In several diseases, the BMDD deviates from this reference because of differences in remodelling and/or mineralisation processes [61]. In cortical bone, the mean mineral content depends on skeletal site [36], so that cortical BMDDs may differ from the reference trabecular BMDD in health too.

Novel measurements of cortical BMDDs from human femur midshafts in healthy adults reveal a higher degree of mineralisation than the reference trabecular BMDD, but a similar spread of mineral heterogeneity around the most frequently occurring calcium content (position of the peak of the BMDD) [11]. To understand the reason for the difference in mineral distribution between trabecular bone and femoral cortical bone in healthy adults, we propose in this paper a systematic hypothesis-testing analysis of BMDDs using an established mathematical model of bone mineral heterogeneity due to remodelling and mineralisation. This mathematical model allows us to quantify BMDD signatures in terms of descriptive parameters of remodelling and mineralisation processes [62, 63].

Shifts of BMDDs and of average bone mineral densities towards higher and lower mineral densities have previously been attributed to lower and higher birth rates of basic multicellular units (BMUs), respectively [6, 46], and acceleration and deceleration of mineral accumulation in specific bone diseases [50]. Mineralisation kinetics, i.e. the accumulation of mineral density with time in a small volume of newly formed bone, has been reported to be similar in cortical and trabecular bones in healthy dogs [41], and in the iliac crest of healthy ewes [3]. Since turnover rates are normally lower in femoral cortical bone than in trabecular bone [40, 53], this difference could explain the shift in BMDD towards higher mineral densities seen in human femoral cortical bone. A lower turnover rate provides more time for secondary mineralisation to take place, which increases the average bone mineral density [6]. However, lower turnover rates also lead to broader BMDD peaks [61, 63, 11]. For the femoral cortical BMDD to possess a spread around the mean similar to that of the trabecular BMDD, additional processes than turnover rate may need to be accounted for. We thus explore systematically and quantitatively further hypotheses related to differences in resorption patterns of remodelling, and to the timescale of mineralisation kinetics. One of the novelties of the mathematical analysis presented in this paper is to investigate whether resorption behaviour of osteoclasts targeting specific calcium content could explain the differences between the BMDDs. Resorption patterns are likely to be different in cortical bone, where resorption may reach any mineral density by tunnelling of new Haversian canals, compared to trabecular bone, where resorption predominantly reaches weakly mineralised bone at the trabeculae’s surfaces [43, 25].

## 2 Materials and Methods

### 2.1 Backscattered electron imaging and experimental BMDD

Midshaft femur cross-sections were extracted post mortem at the Department of Forensic Medicine of the Medical University of Vienna from four adult women (48, 50, 55 and 56 years old), who died unexpectedly with no known bone pathologies. The study was performed in accordance with the ethic commission board of the institution (EK#: 1757/2013). A single femur cross-section was extracted from each donor. The samples were frozen at −20°C immediately after extraction. Prior to measurements, the samples were unfrozen, dehydrated and embedded in polymethylmethacrylate (PMMA) as described elsewhere [57]. Using the scanning electron microscope (Zeiss supra 40, Oberkochen, Germany) in backscattered mode, qBEI scans of in average of 40 mm^2^ bone area per sample covering medial or lateral cortical bone were acquired with a pixel resolution of 1.73 µm. The imaging experimental setup was calibrated and checked for measurement reliability to convert grey intensity values into calcium weight percent (wt% Ca), and further to extract BMDDs following standard procedures [57, 39]. The evaluation was done twice, first by considering the whole cortex and then, by excluding the endosteal and periosteal regions. Obtained differences were minor and, therefore, only results from a whole cortex evaluation are presented. A mean femoral cortical BMDD (Fm.BMDD) was constructed from the very similar individual BMDDs by calculating the mean values for each histogram bin [11].

The reference trabecular BMDD (Tb.BMDD) used in this paper corresponds to the average of 52 BMDDs coming from different individuals, and different skeletal sites from a previous study [58]. Although the number of individuals investigated here to define (Fm.BMDD) is lower than the cohort used to establish the reference trabecular BMDD, the cortical BMDD statistics is substantially improved by the large areas measured. In [58], 5–6 regions of tissue area 5 mm^2^ were used per cancellous bone sample, but the analysed bone area is reduced due to the much smaller BV/TV of trabecular bone compared to femoral cortical bone. In the femur, the degree of mineralisation is known to depend slightly on the region of interest along the periosteal-endosteal axis [64]. However, the variation in BMDD within the cross-sections was small enough to justify the consideration of whole cross-section BMDDs.

The Tb.BMDD and Fm.BMDD were smoothed by the ‘loess’ Matlab procedure [44] with a span of 1.05 wt% Ca. This smoothing enabled better numerical evaluations in the mathematical analyses by alleviating the poor signal-to-noise ratio of the BMDD at very low and very high calcium content [39].

The trabecular and femoral cortical BMDDs exhibit an overall bell-shaped curve (Figure 1). To quantify this shape, we calculated the following shape parameters introduced in Refs [58, 61]: Ca_PEAK_, the calcium content at the peak of the BMDD; Ca_MEAN_, the mean of the BMDD, which corresponds to the average calcium content of the bone sample; and Ca_WIDTH_, the full width at half maximum of the BMDD, which characterises the heterogeneity in mineral density of the sample around the distribution peak [58, 61]. We also calculated the skewness Ca_SKEW_ and the kurtosis Ca_KURT_ of the distribution, which characterise the asymmetry and the peakedness of the distribution, respectively. Skewness and kurtosis are calculated on a restricted interval where the BMDD is larger than 1% of the peak height.

**Figure 1.**
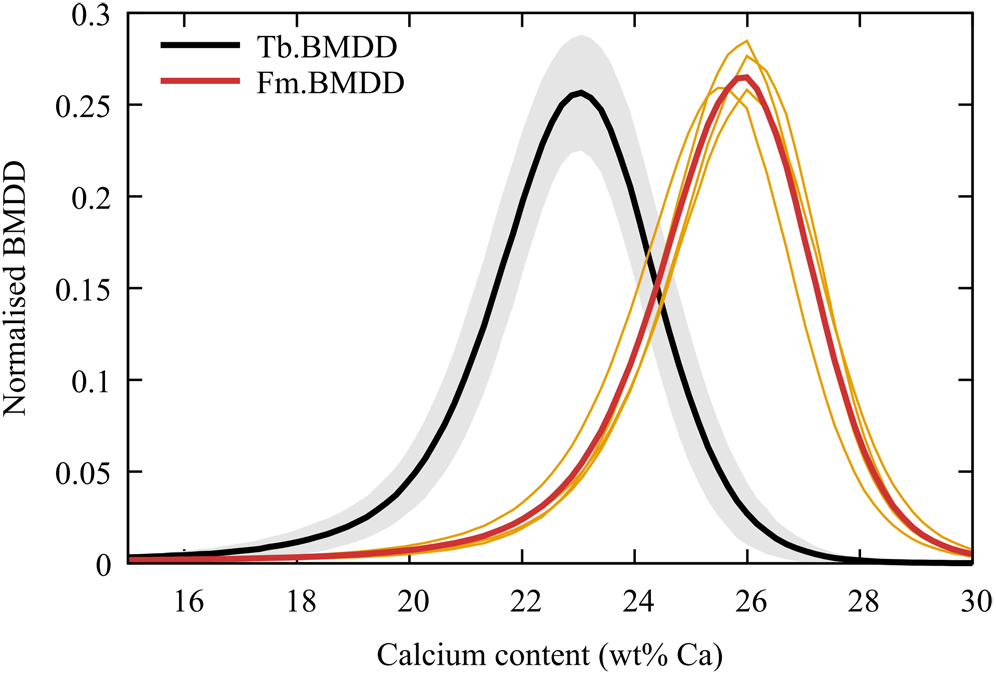
Experimental BMDDs: Reference trabecular BMDD (Tb.BMDD, thick black line) with the ± SD interval (grey lines); adult femoral cortical mean BMDD (Fm.BMDD: thick orange line, with corresponding individual BMDDs as thin orange lines). The BMDDs are normalised to the evaluated bone area (100%).

The strength of the BMDD analysis presented in this study, is to address rather late stages of mineralisation, which are difficult to investigate with labelling techniques.

### 2.2 Mathematical model of the BMDD

To evaluate the influence of turnover rate and mineralisation kinetics on BMDDs, we use the mathematical model developed by [62, 63]. This mathematical model is based on the general, mechanistic principles of the balance of elementary mineralised tissue volumes in a region of interest of bone tissue (TV) during the evolution of the tissue under remodelling and mineralisation processes. The model applies to a region of bone tissue undergoing remodelling, in which mineral density evolves by the processes of (1) new bone formation; (2) bone mineralisation; and (3) bone resorption. Our main assumption is that the mineralisation kinetics is uniform within TV, i.e., independent of space. The influence of these fundamental processes on the BMDD is summarised in Figure 2. We present the main features of this mathematical model below and refer the reader to A for further details. In the mathematical equations, the BMDD is denoted by *ρ*(*c*), where *c* is the calcium content. The quantity *ρ*(*c*)d*c* represents the volume of bone within the sample that possesses a calcium content in the histogram bin [*c*, *c* + d*c*). The remodelling and mineralisation processes are fully characterised in the model by three elements: the mineralisation rate, the resorption rate, and the formation rate, as described below.

**Figure 2.**
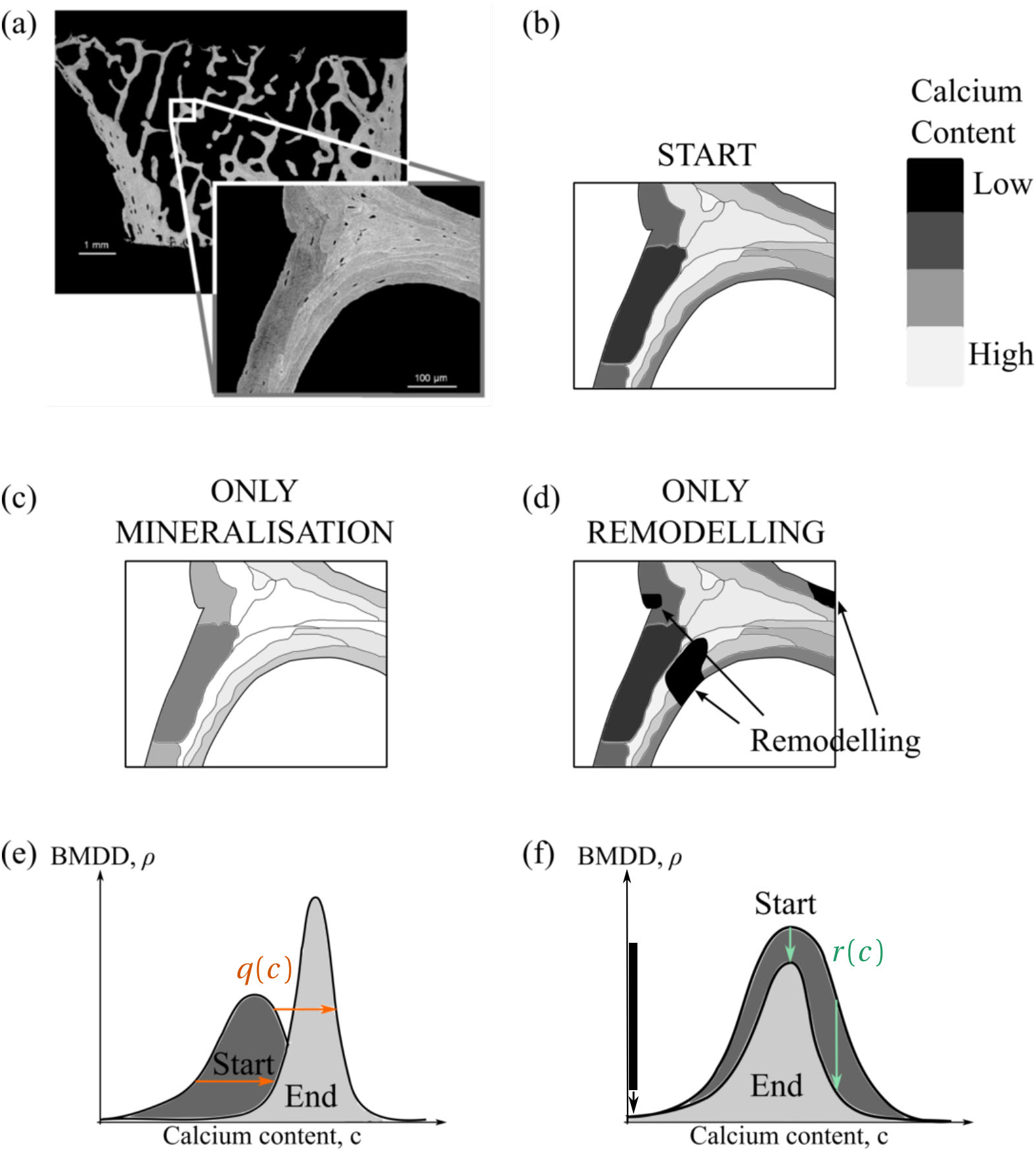
(a) qBEI image of iliac crest trabecular bone. Distinct bone packets within trabeculae are seen as regions of distinct grey colours. (b) Schematic representation of the zoomed qBEI image, where the different degree of mineralisation between the bone packets are highlighted. The influence of mineralisation without remodelling, and of remodelling without mineralisation on the calcium distribution in this region of bone is schematised in (c) and (d), respectively. The corresponding changes in BMDD due to these processes are shown in (e)–(f): (e) mineralisation rate *q*(*c*) (orange arrows) pushes the BMDD towards higher calcium content; the peak width narrows due to the slowdown of mineralisation kinetics as mineralisation proceeds [11]; and (f) bone resorption rate *r* (*c*) (green arrows) removes bone and thus lowers the BMDD while bone formation (black bar) introduces unmineralised bone at *c* = 0.

The mineralisation rate, denoted by *q*(*c*) (in wt% Ca per unit time), describes how fast calcium accumulates in a microscopic region of bone of calcium content *c*. This function is defined such that during a small time increment *Δt*, calcium content increases by *Δc* = *q* (*c*)*Δt*. The mineralisation rate *q*(*c*) entirely determines the kinetics of mineralisation *c* (*t*), i.e. how the calcium content of this region accumulates with time. Conversely, the mineralisation kinetics *c*(*t*) entirely determines the mineralisation rate *q*(*c*), see Eq. (10) [62]. The effect of mineralisation, which occurs at different speeds *q* (*c*) within a sample because of the heterogeneity of mineral density, is to push the BMDD towards higher calcium content, as illustrated in Figure 2(e).

The resorption rate *r* (*c*) is the probability per unit time that bone matrix with calcium content *c* is resorbed by osteoclasts. Different osteoclast resorption patterns can be captured by modifying this function. The effect of this function in the mathematical model is to lower the height of the BMDD with different propensities depending on *c*, as illustrated in Fig. 2(f). A calcium-independent resorption rate represents random resorption, i.e. each region of bone is as likely to be resorbed, irrespective of its calcium content. The quantity *r*(*c*)*ρ*(*c*)d*c* is the volume of bone with calcium content in [*c*, *c* + d*c*) resorbed per unit time. Summing *r* (*c*)*ρ*(*c*)d*c* over all the histogram bins gives the total volume of bone resorbed per unit time in the sample, see Eq. (6).

Finally, the formation rate is the total volume of bone formed per unit time by the osteoblasts. Since osteoblasts form unmineralised matrix, the calcium content deposited by the osteoblasts is taken to be 0 wt% Ca.

Because we investigate differences in BMDDs in healthy adults, we assume that bone resorption and bone formation are balanced. The bone volume formed per unit time and the bone volume resorbed per unit time are taken to be identical, and we express these quantities by the turnover rate *χ*, defined as the fraction of bone volume (BV) renewed per unit time (Eq. (6)) [24]. In humans, turnover rate depends on bone type and skeletal site. In this study, we take the turnover rate in trabecular bone to be *χ*_Tb_ = 20%/year, and the turnover rate in femoral cortical bone to be *χ*_Fm_ = 5%/year [26, 52, 53]. The functions and parameters used in this model are listed in Table 1.

**Table 1.** Nomenclature

| Symbol | Description | Value/Unit |
| --- | --- | --- |
| $c$ | Calcium content | wt% Ca |
| $\rho(c)$ | Bone Mineral Density Distribution (Tb.BMDD; Fm.BMDD) | mm<sup>3</sup>/wt% Ca |
| $q(c)$ | Mineralisation rate | wt% Ca/year |
| $r(c)$ | Resorption function | year<sup>-1</sup> |
| $c(t)$ | Mineralisation kinetics | wt% Ca |
| $\chi_{Tb}$ | Turnover rate in trabecular bone | 20%/year |
| $\chi_{Fm}$ | Turnover rate in femoral cortical bone | 5%/year |

The conservation of bone volume leads to a steady state of the BMDD, due to an equilibrium between (i) the continuous generation of new unmineralised bone due to formation; (ii) the continuous drift of bone towards higher calcium content due to mineralisation; and (iii) the continuous lowering of the BMDD due to resorption (Fig. 2). In steady state, the mathematical model expresses a relationship between the BMDD *ρ*(*c*), the mineralisation rate *q*(*c*), the resorption rate *r* (*c*) and the turnover rate *χ* [62]. Both mineralisation rate and resorption rate combine to make up a BMDD signature (see Eq. (7)). To extract the mineralisation rate *q*(*c*) from measurements of the BMDD (Eq. (8)), assumptions need to be made on the resorption rate *r*(*c*). Equivalently, to extract the resorption rate *r*(*c*) from measurements of the BMDD (Eq. (9)), assumptions need to be made on the mineralisation rate *q*(*c*). To explain the differences observed between Fm.BMDD and Tb.BMDD, we thus test a series of assumptions on differences that may exist in turnover rate, resorption rate, and mineralisation rate between these two bone types.

## 3 Results

The qBEI measurements show that the mean femoral cortical BMDD (Fm.BMDD) is of similar shape as the reference trabecular BMDD of healthy adults (Tb.BMDD), but it is shifted towards higher calcium contents (Fig. 1). This observation is described quantitatively by our measurements of peak shape and location. Table 2 shows that the full width at half maximum Ca_WIDTH_, skewness Ca_SKEW_, and kurtosis Ca_KURT_ are nearly identical for Fm.BMDD and Tb.BMDD. The difference in value of these parameters is well within the standard deviations of the measurements. However, the peak location Ca_PEAK_ of Fm.BMDD is shifted by about 3wt% Ca, well beyond the standard deviations of Ca_PEAK_.

**Table 2.** Shape parameters of the femoral cortical BMDD from adults, the reference trabecular BMDD, and their difference. The trabecular values are taken from Ref. [61].

| BMDD-Parameter | Cortical<br>Mean Values<br>± StD | Trabecular<br>Mean Values<br>± StD | Difference<br>(Cortical – Trabecular) |
| --- | --- | --- | --- |
| $Ca_{PEAK}$ [wt% Ca] | 25.95 ± 0.22 | 22.94 ± 0.39 | +3.01 |
| $Ca_{MEAN}$ [wt% Ca] | 25.05 ± 0.28 | 22.20 ± 0.45 | +2.85 |
| $Ca_{WIDTH}$ [wt% Ca] | 3.20 ± 0.10 | 3.35 ± 0.34 | −0.15 |
| $Ca_{SKEW}$ [−] | −0.91 ± 0.21 | −0.90 ± 0.41 | −0.01 |
| $Ca_{KURT}$ [−] | 5.16 ± 0.68 | 5.17 ± 1.31 | −0.01 |

In the following, we explore computationally a series of hypotheses on the intrinsic remodelling and mineralisation processes of femoral cortical and trabecular bone with the aim to understand how the mineral heterogeneity in these bone types can be distributed similarly, but around different peak positions.

### Hypothesis 1

*The difference between femoral cortical and reference trabecular BMDDs is only due to the difference in turnover rate between these two bone types.*

The motivation for this hypothesis is the well-known difference in turnover between bone types [53], and the suspected influence that turnover exerts on the BMDD [6, 63]. As a first step, we therefore assume that

1. Resorption rate is slower in cortical bone than in trabecular bone due to the difference in turnover rate, i.e., the total volume fraction resorbed per unit time is lower in cortical bone;
2. Resorption rate is unbiased in the two bone types, i.e., osteoclasts resorb bone independently of calcium content (random resorption);
3. Mineralisation rate is identical in the two bone types.

To test Hypothesis 1, we check whether Fm.BMDD and Tb.BMDD measured experimentally are compatible with BMDDs calculated from Eq. (7) with the three assumptions above. When resorption is random, the resorption rate functions *r* (*c*) are constant and equal to the turnover rates *χ*_Fm_ or *χ*_Tb_, respectively (see Eqs (5)–(6)). Two possible choices can then be made to determine the common mineralisation rate *q*(*c*) from the experimental data. The first choice corresponds to the mineralisation rate determined from Tb.BMDD and *χ*_Tb_ using Eq. (8), as was done in Ref. [62]. The second choice corresponds to the mineralisation rate determined from Fm.BMDD and *χ*_Fm_ using Eq. (8). By means of Eq. (7), Hypothesis 1 then allows us to estimate computationally Fm.BMDD from the experimental Tb.BMDD, denoted by “Fm.BMDD (Hyp.1)” in Figure 3, and to estimate computationally Tb.BMDD from the experimental Fm.BMDD, denoted by “Tb.BMDD (Hyp.1)” in Figure 3. Figure 3 shows that the BMDDs computed using either of these choices of mineralisation rates differ significantly from the BMDDs measured experimentally. In particular, Fm.BMDD (Hyp.1) has a Ca_PEAK_ of 24.5 wt% Ca and a Ca_WIDTH_ of 5.15 wt% Ca, and Tb.BMDD (Hyp.1) has a Ca_PEAK_ of 24.5 wt% Ca and a Ca_WIDTH_ of 2.80 wt% Ca (compare with the experimental values in Table 2).

**Figure 3.**
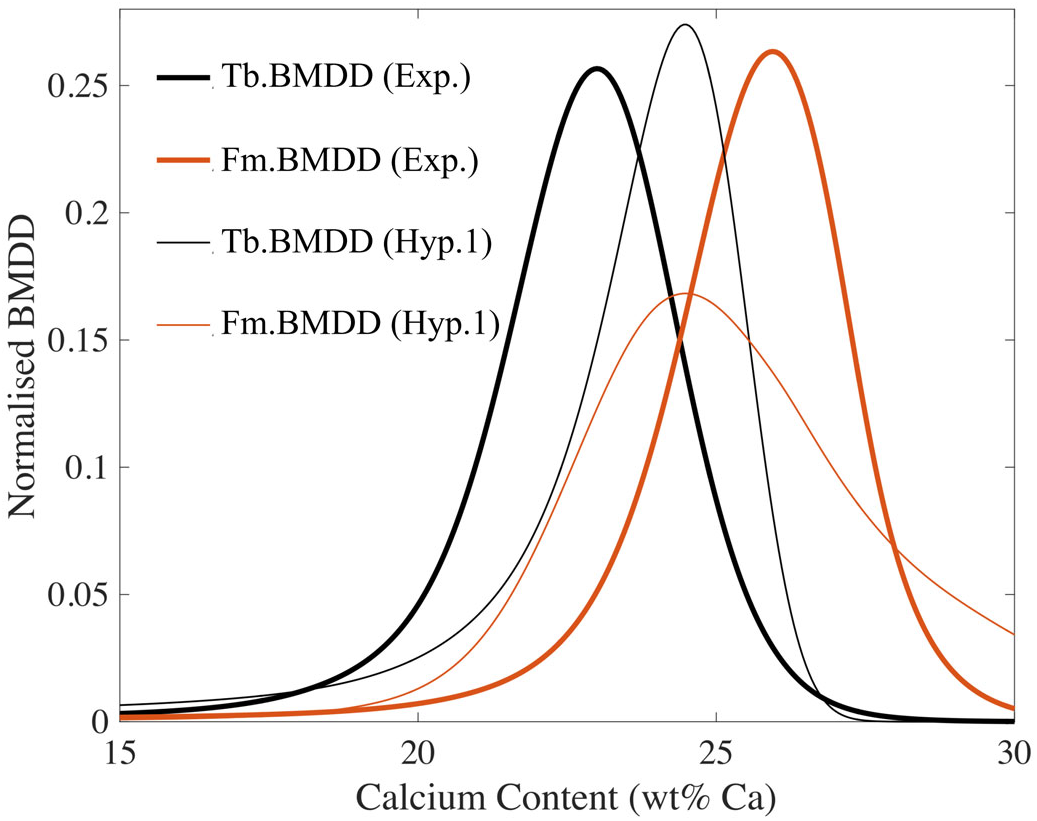
Comparison between experimental BMDDs (thick lines) and BMDDs computed with Hypothesis 1 (thin lines). There is a large discrepancy between the measured femoral cortical BMDD (thick orange) and the BMDD computed with cortical turnover rate and mineralisation rate extracted from the reference trabecular BMDD (thin orange). There is also a large discrepancy between the reference trabecular BMDD (thick black) and the BMDD computed with trabecular turnover rate and mineralisation rate extracted from the femoral corticalBMDD (thin black).

The mathematical model, therefore, shows that turnover rate influences Ca_PEAK_ as expected, i.e., Ca_PEAK_ is low at high turnover rate and Ca_PEAK_ is high at low turnover rate. But turnover rate has also a strong influence on peak width Ca_WIDTH_. Both experimental BMDDs have a similar Ca_WIDTH_ (Table 2), so that no value of turnover rate difference between femoral cortical bone and trabecular bone is able to match jointly Ca_PEAK_ and Ca_WIDTH_ of the computed BMDDs with those of the measured BMDDs [62, 11]. Thus, the difference in mineral distribution between the two bone types cannot be attributed to a change in turnover rate alone.

### Hypothesis 2

*The difference between femoral cortical and trabecular BMDDs is due to a difference in turnover rate, and to an osteoclastic bone resorption biased towards specific calcium contents in* one *of the two bone types.*

The motivation for this hypothesis is the likely difference in resorption pattern between femoral cortical bone and trabecular bone. As a second step in our computational hypothesis testing, we still assume random resorption in trabecular bone, but explore whether there are biased resorption patterns in femoral cortical bone such that the BMDD computed from Eq. (7) matches the measured femoral cortical BMDD. Biased resorption toward specific calcium contents may represent a combination of mineral-density-dependent speed of resorption, and preferential initiation of resorption in regions or surfaces of bone with specific mineral densities. Mathematically, these situations are captured by letting the resorption rate be a general function *r*(*c*) of calcium content. The unique resorption function *r*(*c*) that is consistent with the measured femoral cortical BMDD can be computed using Eq. (9). Figure 4 shows that this function is negative for a wide range of values of calcium content. Conversely, assuming random resorption in cortical bone, the unique trabecular bone resorption function such that the BMDD computed from Eq. (7) matches the measured reference trabecular BMDD is also negative for a range of values of calcium content (Fig. 4). Negative parts of *r*(*c*) for *c >* 0 represent the formation of readily mineralised bone. The area under the negative region of the graphs of *r*(*c*)*ρ*(*c*)*/χ* shown in Figure 4 represents the total volume of bone that would be formed readily mineralised per turnover time. Under Hypothesis 2, a significant amount of bone would be formed readily mineralised, which clearly contradicts what is known about osteoblast action and bone mineralisation. Hypothesis 2 is thereby falsified.

**Figure 4.**
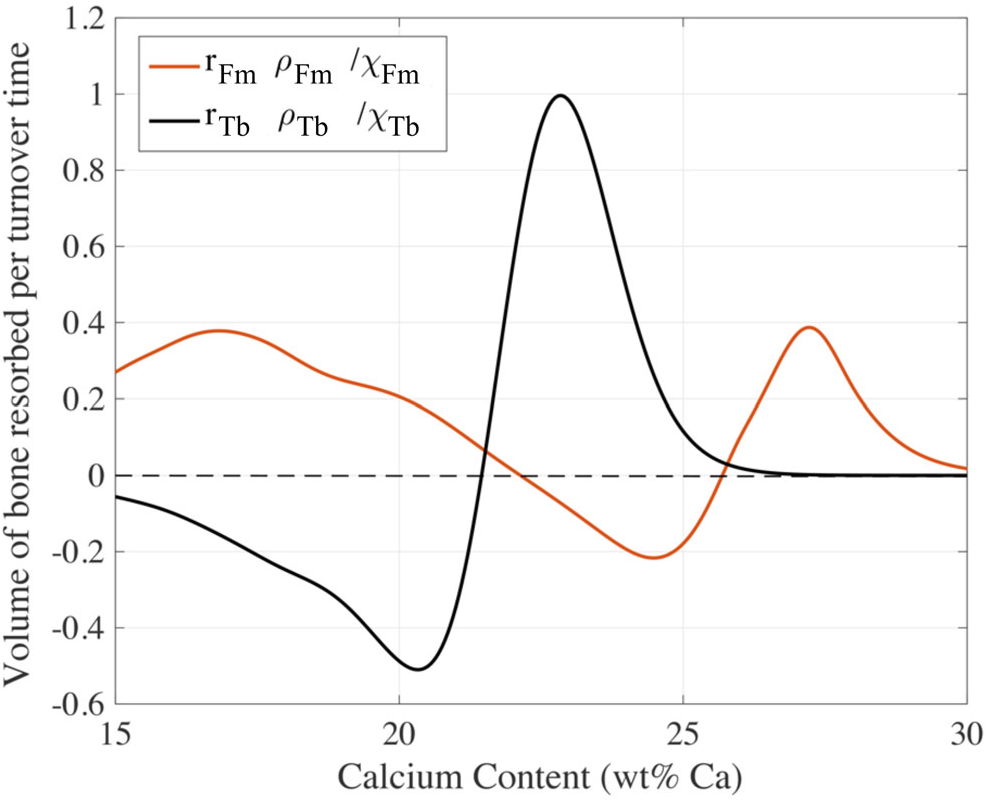
Distribution of volume of bone resorbed per unit time under Hypothesis 2 with random resorption assumed in trabecular bone (orange), and random resorption assumed in femoral cortical bone (black). The unique resorption rate functions *r*(*c*) for which Hypothesis 2 holds are negative in a region of calcium content, which means Hypothesis 2 would only hold if there is formation by osteoblasts of readily mineralised bone.

### Hypothesis 3

*The difference between femoral cortical and trabecular BMDDs is due to a difference in turnover rate, and to osteoclastic bone resorption biased towards specific calcium content differently in* both *bone types.*

As a third step, we now relax the assumption that random resorption occurs in one of the bone types and we attempt to determine whether there are two general resorption rate functions *r*_Fm_(*c*) and *r*_Tb_(*c*), such that the BMDDs computed from Eq. (7) match the experimental BMDDs. Mineralisation kinetics is still assumed to be identical in both bone types as in Hypotheses 1 and 2, i.e., we still consider Assumption (3) to hold. The analysis is now complicated by the fact that mineralisation rates are determined from the experimental BMDDs under a choice of resorption functions by Eq. (8),. The procedure employed to test Hypothesis 3, is therefore (i) to first choose a resorption function *r*_Tb_(*c*); (ii) to deduce the mineralisation rate from Eq. (8) using the femoral cortical BMDD and the choice of resorption function *r*_Tb_(*c*); and (iii) to then determine *r*_Fm_(*c*) from Eq. (9) using the reference trabecular BMDD. We can also proceed conversely by choosing first *r*_Fm_(*c*), and calculate how this choice determines *r*_Tb_(*c*). This procedure is repeated to explore the set of all possible choices of positive resorption functions *r*_Tb_(*c*) that lead to positive resorption functions *r*_Fm_(*c*), and vice versa. Such a pair of positive resorption functions *r*_Tb_(*c*), *r*_Fm_(*c*) represents targeted osteoclast resorption behaviours that lead to the experimental BMDDs while having identical mineralisation kinetics in femoral cortical bone and trabecular bone. Below, we show that this set is strongly restricted by the requirement that both resorption functions must be positive. We find that only cortical resorption functions that strongly target very lowly mineralised bone satisfy this requirement.

The detailed mathematical analysis is more involved than in the previous cases, since it requires the exploration of a large space of possible resorption functions. The reader is referred to B for the mathematical details. To help us enforce the positivity of *r*_Fm_(*c*) given a choice of *r*_Tb_(*c*), we introduce an auxiliary function *G*_Tb_(*c*) defined in Eq. (14). Choosing an initial resorption function *r*_Tb_(*c*) is equivalent to choosing an auxiliary function *G*_Tb_(*c*). The advantage is that to explore the set of all resorption functions *r*_Tb_(*c*) that give rise to positive resorption functions *r*_Fm_(*c*), it is sufficient to explore the set of all auxiliary functions *G*_Tb_(*c*) below an explicit upper limit determined by the experimental BMDDs (Eq. (17)). Conversely, to explore the set of all resorption functions *r*_Fm_(*c*) that give rise to positive resorption functions *r*_Tb_(*c*), it is sufficient to explore the set of all auxiliary functions *G*_Fm_(*c*) below an explicit upper bound.

To illustrate the difficulty of enforcing the positivity of the resorption functions, Figures 5a,c show a number of different initial choices of *r*_Fm_(*c*) that correspond to osteoclasts resorbing cortical bone that is preferentially (i) lowly mineralised, by assuming that *r*(*c*) = *r*_0_ + *λ*(e^*γc*^ − 1) with negative exponents *γ* = −10, −30 and *r*_0_ = 5.72 10^6^, 15, respectively; (ii) highly mineralised by assuming that *r*(*c*) = *r*_0_ + *λ*(e^*γc*^ − 1) with positive exponents *γ* = 5, 20 and *r*_0_ = 0.025; and (iii) of calcium content restricted to Gaussians of different centres *M* and widths *σ*, i.e., 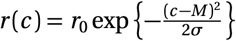. Since the resorption functions must satisfy the turnover condition (6), they only depend on two independent parameters. Figures 5b,d show that choosing functions *r*_Fm_(*c*) that correspond to preferential resorption targeting intermediate or highly mineralised cortical bone (*γ* = 5, 20, and *M* = 25, 28) do not lead to trabecular resorption rate functions *r*_Tb_(*c*) that are positive. The auxiliary functions *G*_Fm_(*c*) corresponding to these choices of *r*_Fm_(*c*) lie in part above the upper limit (Fig. 5b,d). The same holds for a mild resorption preference towards low calcium content (*γ* = −10). The only resorption functions *r*_Fm_(*c*) that lead to positive trabecular resorption functions *r*_Tb_(*c*) are the Gaussian centred around lowly mineralised bone (*M* = 15), and the exponential resorbing only lowly mineralised bone (*γ* = −30). The procedure to check Hypothesis 3 thoroughly is to explore the full set of initial choices *G*_Tb_(*c*) below the upper limit and inspect the cortical resorption functions *r*_Fm_(*c*) determined by these choices (see B). For illustrative purposes, we show in Fig. 6 the distribution of cortical bone volume resorbed per unit time *r*_Fm_(*c*)*ρ*Fm(*c*) when *G*_Tb_(*c*) is set as the upper limit (borderline case, Eq. (17)). In this case too, cortical bone would be preferentially resorbed at extremely low calcium content (< 22wt% Ca).

**Figure 5.**
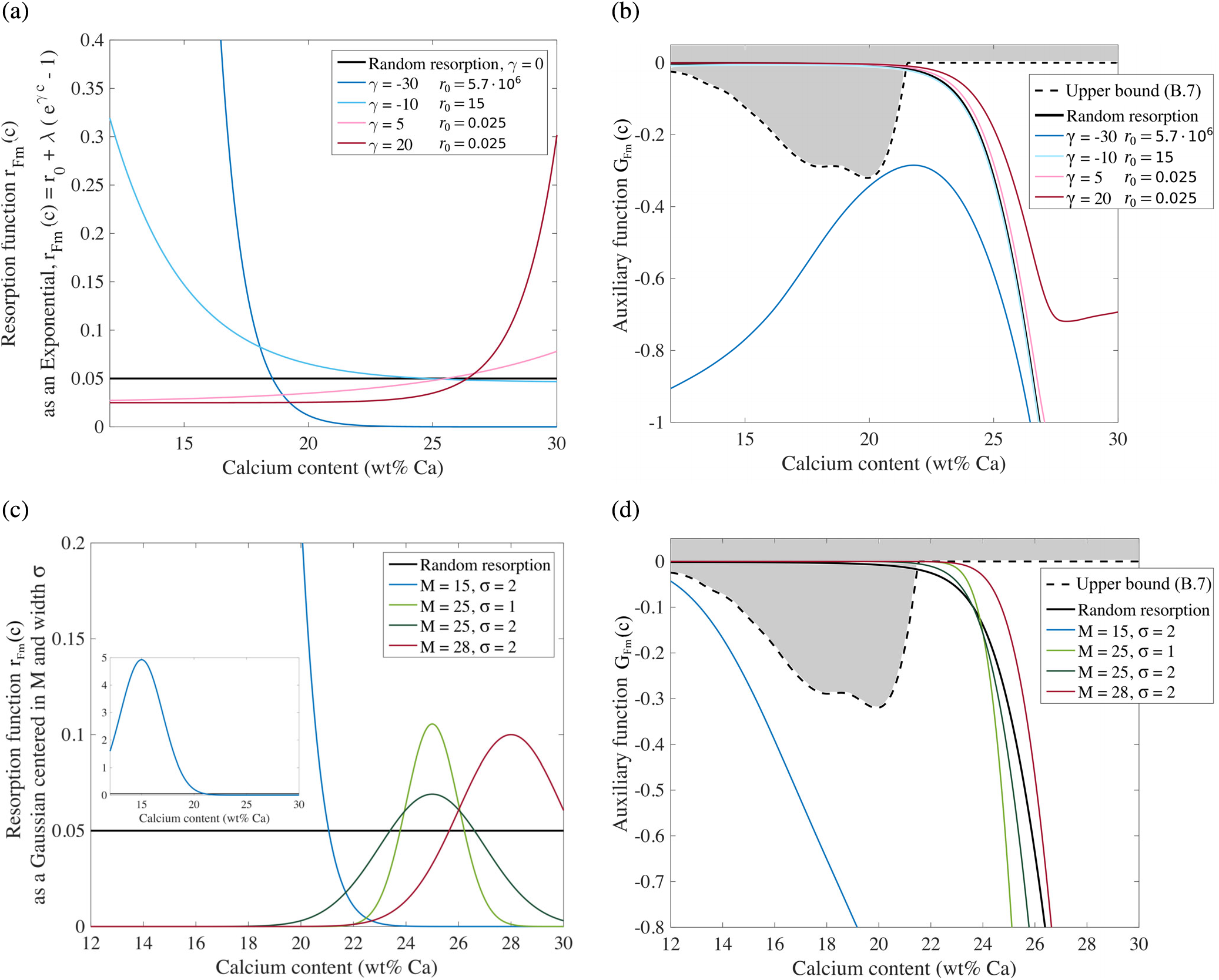
Resorption functions used to test Hypothesis 3; (a) exponential resorption functions; and (c) Gaussian resorption functions. The inset in (c) shows a broader view of the Gaussian centred at 15wt% Ca. The auxiliary functions *G*_Fm_(*c*) corresponding to these exponential and Gaussian resorption functions are shown in (b) and (d), respectively. Auxiliary functions *G*_Fm_(*c*) are subject to the condition (17) derived in B, i.e., they must sit below the shaded area bounded by the dashed black line to ensure that resorption functions are positive, i.e., that no bone is formed readily mineralised. Only resorption functions strongly targeting lowly mineralised bone satisfy this constraint.

**Figure 6.**
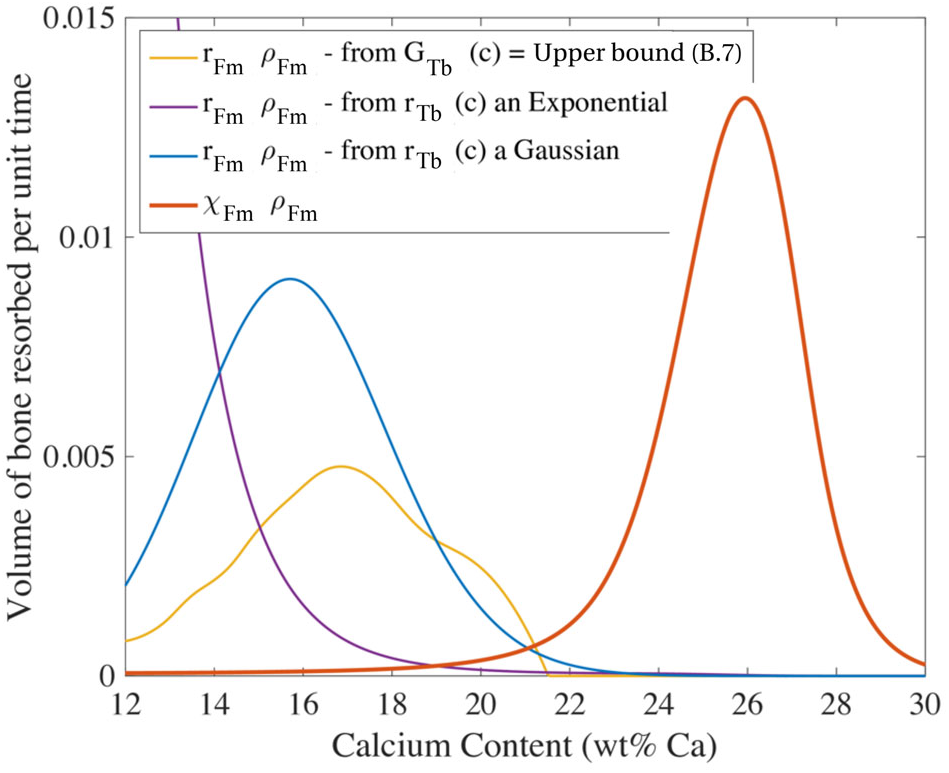
Distribution of volume of bone resorbed per unit time in cortical bone, *r*_Fm_(*c*)*ρ*Fm(*c*), obtained for certain choices of trabecular auxiliary functions *G*_Tb_(*c*), including for when *G*_Tb_(*c*) is set as the upper bound of the condition (17). The measured femoral cortical BMDD (rescaled by *χ*_Fm_) is also shown to emphasise that the cortical resorption behaviours obtained in these cases renew a very small volume of lowly mineralised cortical bone at an extremely high rate.

In B.1, we show rigorously that under Hypothesis 3, for any choice of auxiliary function, at least 25% of the femoral cortical bone resorbed per unit time would have a calcium content less than 21wt% Ca, and at least 15% would have a calcium content less than 20 wt% Ca (see Eq. (21) and Fig. 10). In other words, more than a quarter of all the cortical bone renewed would be targeting new, poorly mineralised bone, to the extent that more than 15% of all cortical bone would be resorbed before undergoing secondary mineralisation. These predictions clearly conflict with the concept that bone remodelling is a repair mechanism that enables bones to maintain structural integrity despite fatigue loading [42, 14, 5], especially in mechanically stimulated bone such as the femoral cortex. For all the above reasons, we therefore deem Hypothesis 3 to be falsified.

### Hypothesis 4

*The difference between femoral cortical and trabecular BMDDs is due to a difference in turnover rate, and a difference in mineralisation kinetics between the two bone types.*

Having exhausted all the possibilities in which mineralisation kinetics was assumed identical between trabecular and femoral cortical bone, we now hypothesise that mineralisation kinetics may be different in trabecular bone and femoral cortical bone, i.e., we no longer consider our assumption (3) to hold. To explore this hypothesis computationally, we revert to the assumption of random resorption in both bone types for simplicity. In this case, the mineralisation rates *q*(*c*) and mineralisation kinetic laws *c*(*t*) can be determined in each bone types independently by Eqs. (8) and (10) from the experimental BMDDs, Tb.BMDD and Fm.BMDD. The mineralisation rates and kinetic laws determined in this way are shown in Figure 7. Both mineralisation kinetics display a fast initial increase in calcium content, followed by a slower increase in calcium content with time. However, the overall rate of mineralisation in femoral cortical bone is slower than the overall rate of mineralisation in trabecular bone. The timescale of mineralisation kinetics in either bone type depends on the assumed turnover rate in Eq. (8). To factor out the influence of the choice of turnover rate and compare qualitatively the mineralisation kinetics in femoral cortical bone and trabecular bone, we perform a scaling in time of the mineralisation kinetics of cortical bone by the ratio of turnover rates 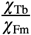. Remarkably, Figure 7 shows that the time-scaled cortical mineralisation rate and time-scaled cortical mineralisation kinetic laws have similar shape as the trabecular mineralisation rate and mineralisation kinetic law, respectively, but are shifted along the calcium content axis. Accounting for this shift additionally by including an offset *κ* results in a nearly perfect superposition of the transformed cortical mineralisation functions and the trabecular mineralisation functions, i.e.,

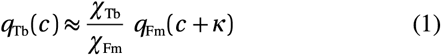

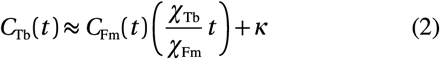

**Figure 7.**
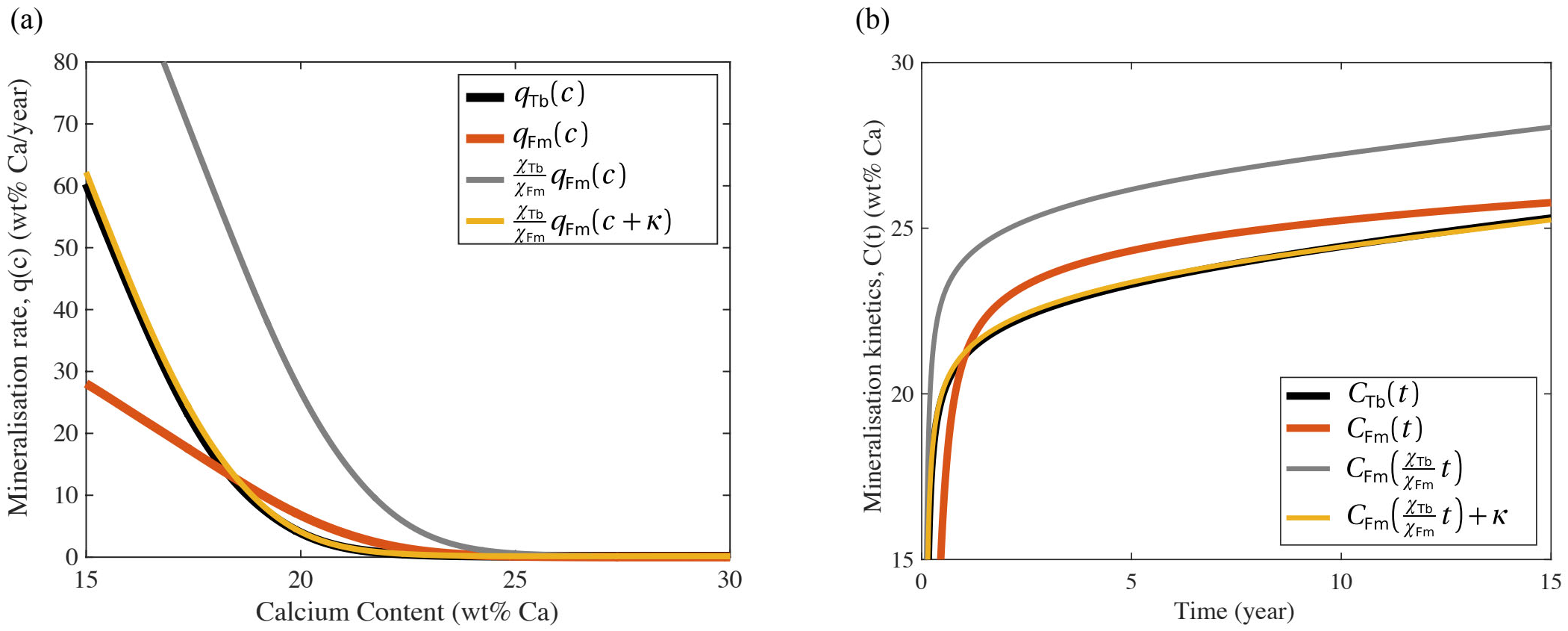
(a) Mineralisation rates *q*_Tb_(*c*), *q*_Fm_(*c*); and (b) Mineralisation kinetic laws *c*_Tb_(*t*), *c*_Fm_(*c*) obtained from the experimental trabecular BMDD and femoral cortical BMDD under Hypothesis 4. The mineralisation rates *q*(*c*) and kinetic laws *c*(*t*) obtained by rescaling the timescale of femoral cortical mineralisation by the ratio of turnover rate 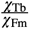, and by shifting the calcium axis by *κ* ≈ 2.83 wt% Ca are also shown. The time-rescaled and calcium-shifted femoral cortical mineralisation functions *q*(*c*) and *c*(*t*) fit the trabecular mineralisation functions *q*_Tb_(*c*) and *c*_TB_(*t*) remarkably well both at low calcium densities (≲ 20 wt% Ca, as seen more obviously in (a)), and at high calcium densities (≳ 20 wt% Ca, as seen more obviously in (b)).

The goodness of match between the mineralisation kinetics of trabecular and femoral cortical bone after a scaling in time and a shift in calcium content in Eqs (1)–(2) is remarkable because it holds in the full range of experimentally accurate measurements of calcium densities (Fig. 7). It is important to note that this match does not rely on the choice of ratio of turnover rates. The same match holds for other assumed values of turnover rates, so long as these values are used consistently in the other quantities that depend on them. It can be shown mathematically from Eq. (7) that the reason for this goodness of match is the similarities of the *shapes* of the two BMDDs (Fig. 1 and Table 2). By translating the femoral cortical BMDD towards lower calcium contents, such that both peak locations coincide, the two BMDDs superimpose almost perfectly. The translated femoral cortical BMDD is within the 95% confidence interval of the trabecular measurements. A sensitivity analysis of the link between BMDD shape and mineralisation kinetics is provided in C.

## 4 Discussion

Turnover rate is well-known to vary across skeletal sites and bone types. In particular, regions of femoral cortical bone of low porosity may be turned over roughly 4 times slower than trabecular bone [26, 52, 53]. This has been proposed to explain the high degree of mineralisation of cortical bone compared to trabecular bone [6]. However, our analysis clearly excludes that the difference in mineral density distribution between femoral cortical bone and trabecular bone captured in the BMDDs is an effect of turnover only (Hypothesis 1, Fig. 3). By testing rigorously general hypotheses on resorption rates, we were able to exclude other possible ways by which one may explain the difference between the two BMDDs. In particular, the concept that different resorption behaviours of osteoclasts may take place in cortical versus trabecular bone, was found to be an insufficient explanation (Hypotheses 2 and 3, Fig. 4–6). The assumption that osteoclasts may resorb preferentially bone with specific calcium content can be understood as (i) a cellular preference to target microcracks, generally situated in old, highly mineralised bone [14, 42]; (ii) geometrical constraints related to the fact that resorption occurs first at existing surfaces in trabecular bone. Bone tissue near the bone surfaces is less mineralised than interstitial bone on average, so resorption targeting less mineralised bone represents resorption of bone layers that remain close to the bone surface. In contrast, resorption targeting more mineralised bone represents the resorption of deeper layers of bone, such as interstitial bone, which is located further away from bone surfaces. Assigning various propensities for bone tissue resorption depending on mineral content effectively captures such fine-scale spatial dependences of bone resorption. Because we make no a priori restriction on the functional relationship between calcium content and resorption rate, we do not need to specify further how mineral density correlates with spatial location within bone to account for these effects; and (iii) mechanical regulation of remodelling, which may recruit osteoclasts at different rates depending on the mechanical properties of the bone site [13] such as the local stiffness of bone matrix, which depends in particular on mineral content. Our completely general choice of possible resorption functions *r*(*c*) thereby takes into consideration in an implicit manner a broad range of influencing factors on osteoclastic behaviour.

The fact that the mineralisation kinetic laws in Figure 7 only match after rescaling in time suggests that mineralisation kinetics is different in trabecular and femoral cortical bone. Eqs (1)–(2) indicate that the time scale of mineralisation is coupled with the time scale of bone turnover independently of bone type (see D). Our conclusion that mineralisation may proceed at a different rate in trabecular and femoral cortical bone focuses on mid/late stages of mineralisation. In a previous study, Marotti et al. studied the mineralisation kinetics by fluorescence labelling over 47 days in the dog humerus, radius, ulna, femur, tibia diaphysis, and radius metaphysis, and found no statistical differences [41]. These findings hold for early stages of mineralisation and are thus not contradictory with our conclusions. At this early stage of mineralisation, our computational model predicts a mineralisation kinetics which is very similar for the two bone types (Fig. 7). In a more recent study, Bala et al. investigated the mineralisation kinetics in iliac crest of ewes over 30 months [3], also finding no statistical differences of mineralisation kinetics between trabecular and cortical bone. This observation emphasises the importance of the skeletal site for bone mineral heterogeneity. Our conclusions are drawn from BMDD data from femoral cortical bone. The development of the cortex in the iliac crest follows an intricate development during growth [29] that, unlike femoral bone, is likely to retain traces of its growth history in terms of large-scale spatial dependences of mineral heterogeneity. Furthermore, its cortical and trabecular compartments may be regulated similarly [45]. Our analysis relies on a steady state assumption where traces of growth history have been erased by remodelling. The mathematical model also relies on the continuum modelling assumption that the tissue volume TV is large enough so that the BMDD averages many mineral heterogeneities, but small enough that there are no large-scale spatial dependences of the BMDD within TV.

The experimental basis of our study is that the peak position of Tb.BMDD and Fm.BMDD is different, but the mineral heterogeneity is similar, as quantified by Ca_WIDTH_ (Table 2). One major intent of the current work is to emphasise that this similar heterogeneity is unexpected from our current understanding of the processes of remodelling and mineralisation. If mineralisation kinetics were identical in trabecular and femoral cortical bone, our model predicts that the lower turnover rate in cortical bone would result in a broadening of the BMDD. This broadening is analogous to the broadening of the age distribution of humans when life expectancy increases: the human population becomes simply redistributed on a larger variety of possible ages. Our experimental observation that peak width in Fm.BMDD and Tb.BMDD is not significantly different seems to be observed in other situations of changed bone turnover analysed in terms of the BMDD:

i. The effect of different bisphosphonates on bone turnover [23] and the BMDD [60] have been extensively studied over the last twenty years. Depending on their potency bisphosphonates can reduce the rate of bone remodelling substantially [30, 49]. After 10 years of alendronate treatment the mineral heterogeneity was not significantly different from the trabecular reference BMDD [59]. In a separate study, 5 years of risedronate treatment resulted in a Ca_WIDTH_ not significantly different from its baseline value [66].
ii. Hypoparathyroidism is a condition of known low bone turnover. In vitamin-D treated hypoparathyroid patients the activation frequency in trabecular bone of iliac crest biopsies was reduced by a factor 4.5 in these patients compared to healthy individuals [37]. While the BMDD in the hypoparathyroid patients was significantly shifted towards higher values, the heterogeneity of mineralisation around the peak was not changed compared to the reference trabecular BMDD [48].
iii. Our study also provides a new perspective on the definition of the trabecular reference BMDD of healthy adults [58]. The reference BMDD is defined based on the average of 52 individual BMDDs, since it was found that in healthy adults the trabecular BMDD does not depend on age, gender, ethnicity and skeletal site. The reference value for Ca_WIDTH_ was reported to be 3.35 wt% Ca with a standard deviation of ±0.34 wt% Ca (Table 2). Translating this standard deviation in a variability of turnover using Fig. 6 in Ref. [62] shows that the corresponding variability in turnover would range from −40% to +10% of the average value, a remarkably narrow range having in mind the natural variability of turnover across bone sites and individuals, which is expected to be much larger [53]. The fact that the trabecular BMDD is so consistent despite large variability of turnover rate suggests that another mechanism compensates the influence of turnover rate on mineral heterogeneity in trabecular bone.
iv. The BMDD of cortical bone measured in transiliac bone samples was found to have a much lower Ca_PEAK_ value [45] compared to the Ca_PEAK_ value of cortical bone at the midshaft femur (Fig. 1). Consequently, a reference cortical BMDD (independent of skeletal site) of healthy adults cannot be defined, in contrast with the reference trabecular BMDD. The similarity of Ca_WIDTH_ in Tb.BMDD and Ca_WIDTH_ in Fm.BMDD is inconsistent with what one would expect if differences in mineral density distributions would result from differences in turnover rate alone. If only turnover rate would differ, our model shows that Fm.BMDD would have a broader peak than Tb.BMDD, which is not what is observed.
v. Data from the control group of the risedronate study in Ref. [66] shows that three years of treatment with calcium and vitamin D supplementation (with unknown consequences on the turnover rate) only resulted in a statistically significant shift of the BMDD peak by 0.73wt% Ca [9, 66, 27]. This shift in peak location, presumably due to the extra calcium intake of the patients, was not associated with a change in BMDD peak width compared to baseline values.

A possible interpretation of our observations that the heterogeneity of mineral content around the mean seems rather uniform, is that the processes of remodelling and mineralisation are interlinked (see Eqs (22)–(23)). Indeed, our model shows that if mineralisation rate scales with turnover rate, then the degree of heterogeneity of mineral density around the mean, measured in particular by Ca_WIDTH_, is independent of turnover rate.

A turnover-limited mineralisation kinetics suggests a limitation imposed by the transport and recycling of minerals. If mineral transport is spatially restricted, minerals that are embedded into bone matrix during bone mineralisation may come mostly from the recycling of minerals freed from the bone matrix during bone resorption [56]. The abundance of these recycled minerals depends on turnover rate.

In the studies (i)-(iv) mentioned above the width of the BMDD peak was virtually unchanged, but the peak position typically shifted significantly. In our study an additional shift along the calcium content axis was needed to match the mineralisation kinetics of trabecular and femoral cortical bone. These shifts in the BMDD peak location suggest the existence of an additional calcium reservoir in bone that may be filled to different levels in trabecular and cortical bone during early stages of mineralisation. Indeed, the shift in calcium density between the mineralisation kinetic laws *c*(*t*) of trabecular bone and of femoral cortical bone is generated during the first 1–2 years of mineralisation, see Fig. 7. Other works have proposed the existence of mineral reservoirs in association with the lacunocanalicular network of the osteocytes [17, 15]. In the context of a revived discussion about osteocytic osteolysis [65, 8], the osteocytes are thought to play an active role in the mineralisation process by modifying the perilacunar and pericanalicular matrix [35, 32]. In a recent study on human osteons, the local Ca content measured by qBEI was spatially correlated with the density of the lacuno-canalicular network measured by confocal microscopy. In line with the hypothesis of a mineral reservoir associated with the canalicular network, regions with a dense network were found to have a higher mineral content [55].

Two alternative explanations for the difference between Tb.BMDD and Fm.BMDD should be mentioned. The first relies on the idea that the amount of Ca that can be incorporated into the bone matrix is limited and that this limitation may reduce Ca_WIDTH_ in low turnover scenarios [46, 45]. Our previous computational analysis [11] exploring how a reduced capacity of mineral uptake influences the BMDD in late stages of mineralisation showed indeed that peak broadening is suppressed in low turnover rate scenarios. However, this computational analysis also predicted an increase in the BMDD’s skewness which is not observed experimentally [11]. The second possible explanation is based on the lack of reliable data about turnover rates depending on different skeletal sites and, in particular, different bone types (i.e., cortical vs. trabecular) [53]. If femoral cortical bone and trabecular bone are turned over with more similar rates than what is usually assumed, then the similarity in the shape of the BMDDs of these different bones is also less surprising.

A strength of the quantitative hypothesis-testing approach used in this work is to be able to investigate the influence of various biological processes in isolation. The mathematical model we based our analysis on uses very general principles of mass conservation laws in which bone is created, matures, and is resorbed. The manner in which these processes occur was subject only to recovering the experimentally measured BMDDs. Nevertheless, the model makes two fundamental assumptions most important for the interpretation of our results. First, it is assumed that the temporal incorporation of minerals in bone matrix follows the same kinetics for bone formed at different location and different times in the same qBEI image. This assumption allows the definition of a mineralisation rate *q*(*c*), which can be different depending on bone type, but is otherwise the same for each newly formed bone packet in this bone type. While osteocytes may interfere with the time course of mineral accumulation during the mineralisation of bone matrix [65, 2, 4, 55], most measurements of mineralisation kinetics so far suggest that mineral accumulation remains a good proxy of tissue age within a same qBEI scan. Second, our analysis of adult BMDDs assumes that these mineral distributions are in steady state, i.e. they would not evolve in time under the given conditions of remodelling. This assumption seems justified considering that the age of the individuals is between 48 and 56 years old. However, this steady-state assumption may have to be revisited when interpreting BMDD data of individuals afflicted recently by certain bone disorders, or data from children due to their growth [29].

The mathematical model presented in this paper relies on the rigorous principles of the balance of elementary mineralised tissue volumes under remodelling and mineralisation processes. We applied these principles by considering a spatial scale that contains a similar level of detail as the experimental data. The model therefore includes mineral heterogeneity, but it does not include further variables such as spatial dependences. More detailed explorations of the influence of space and geometry [12, 38, 1] and the interplay with mechanics for the regulation of mineralised tissues [33, 31, 22, 18] are of high interest for future works. The microarchitecture of trabecular bone gives rise to complex strain patterns, which could influence not only the remodelling, but also the mineralisation process. The consideration of bone tissues that retain traces of the history of their mode of growth in the distribution of mineral density is important to be able to analyse cortical BMDDs from the iliac crest, from children [29], and from diseased states.

In summary, our findings suggest that trabecular and femoral cortical bone are more different than usually assumed. The common view is that both bone types are made up of bone lamellae, but that there is a difference in the spatial arrangement of the lamellae, and a difference in the rate at which the bone is turned over. Our conclusion, drawn from quantitative hypothesis testing, that the mineralisation kinetics is different in these bone types suggests that this common view needs to be supplemented by the notion that minerals are incorporated in the two bone types differently. This conclusion is further corroborated by qBEI measurements of very low turnover rate trabecular bone, which exhibits a distinctively lower value of Ca_PEAK_ than that of femoral cortical bone despite having a similar low turnover rate [48, 46]. A possible reason for why trabecular bone and femoral cortical bone may exhibit differences in mineralisation kinetics is that these bone types have different mechanical requirements [19, 28].

## Acknowledgments

We thank Paul Roschger for stimulating discussions. PRB and RW acknowledge support by Universities Australia (UA) and German Academic Exchange Service (DAAD) for the Australia–Germany Joint Research Cooperation Award (Project “The cellular control of mineral heterogeneity in bone”).

## A Mathematical model

Ruffoni et al. developed a phenomenological model of the time evolution of the BMDD *ρ*(*c*, *t*) at the tissue scale [62, 63]. This model includes: (i) bone formation; (ii) bone resorption; and (iii) bone mineralisation. The time evolution of the BMDD is governed by the reaction–advection equation

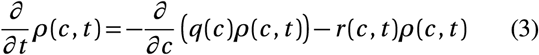

with the boundary condition

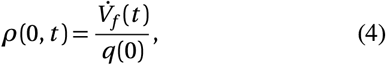

where 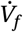 is the total volume of new bone formed per unit time in the sample. In steady state, the total volume of bone formed per unit time equals the total volume of bone resorbed per unit time, and is related to the turnover rate *χ*. Since *ρ*(*c*)d*c* represents the volume of bone with calcium content within [*c*, *c* + d*c*) and *r* (*c*)*ρ*(*c*)d*c* the volume of bone with calcium content within [*c*, *c* + d*c*) resorbed per unit time, one has in steady state:

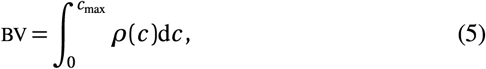

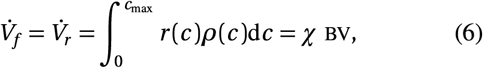

where *c*_max_ is an upper bound for the maximum amount of calcium that can be stored in bone. In a previous study we explored the influence of lower values of the maximum calcium capacity of bone *c*_max_ on the BMDD with the result that in low turnover scenarios, the BMDD peak becomes strongly asymmetric if *c*_max_ is assumed to be smaller than 30 wt% Ca [11]. Here we therefore choose *c*_max_ = 35 wt% Ca.

From Eq. (3), the BMDD in steady state is [62]:

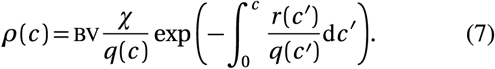

Equation (7) is the central equation used to check whether BMDDs measured experimentally are consistent with the resorption rate functions *r*(*c*) and the mineralisation rate functions *q*(*c*) assumed in Hypotheses 1–4. From Equation (3), we can also obtain the mineralisation rate, *q*(*c*), or the resorption function, *r*(*c*), as a function of the BMDD:

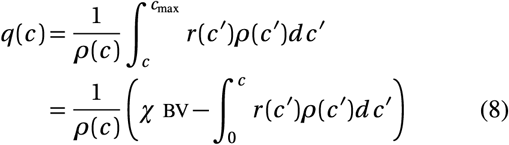

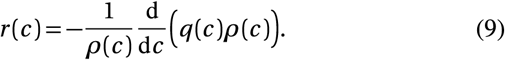

Note that the mineralisation kinetic law *c*(*t*) and the mineralisation rate *q*(*c*) are related by the differential equation

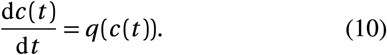

## B Conditions for positive resorption functions in Hypothesis 3

Hypothesis 3 allows both trabecular and cortical resorption functions to depend on calcium content. In this Appendix, we derive necessary and sufficient conditions to ensure that these two resorption functions are positive for all values of calcium content *c*. Since Hypothesis 3 assumes that mineralisation rate is identical in femoral cortical and trabecular bone, i.e. *q*_Fm_(*c*) = *q*_Tb_(*c*), we can express one resorption function as function of the other. Indeed, substituting *q*_Fm_(*c*) occurring in *r*_Fm_(*c*) (Eq. (9)) with *q*_Tb_(*c*) determined from *r*_Tb_ and *ρ*_Tb_ in Eq. (8) gives:

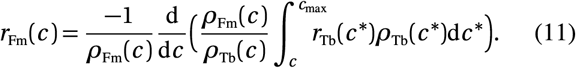

Conversely:

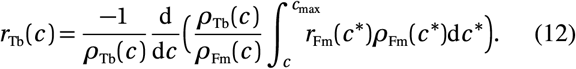

Not all choices of positive resorption functions *r*_Tb_(*c*) in Eq. (11) will lead to a positive resorption function *r*_Fm_(*c*). To ensure that *r*_Fm_(*c*) is positive, *r*_Tb_(*c*) must be such that

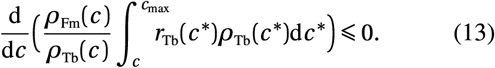

To determine the sign of this derivative, let 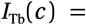 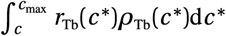 and 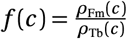 We can rewrite the derivative in Eq. (13) as: 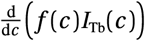. Since both *f*(*c*) and *I*_Tb_(*c*) are positive, we have:

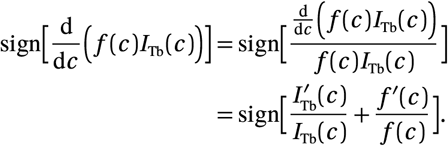

Introducing the auxiliary function:

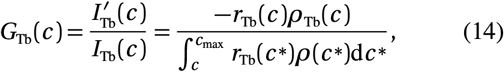

it is clear that the resorption function *r*_Fm_(*c*) is positive if and only if

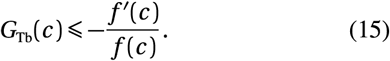

Equation (14) expresses *G*_Tb_(*c*) as a function of *r*_Tb_(*c*). This relationship can be inverted, so that choosing a resorption function *r*_Tb_(*c*) is entirely equivalent to choosing an auxiliary function *G*_Tb_(*c*). Indeed, integrating Eq. (14) over [0, *c*] gives 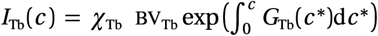. Plugging this expression back into Eq. (14) now expresses *r*_Tb_(*c*) as a function of *G*_Tb_(*c*):

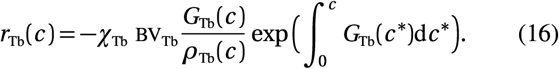

The advantage of introducing the auxiliary function *G*_Tb_(*c*) is that Condition (15) represents a direct restriction on choices of *G*_Tb_(*c*). By definition, *I*_Tb_(*c*) > 0, and 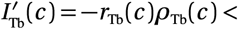 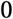, so that *G*_Tb_(*c*) must also be negative. However, the right hand side of In eq. (15) is not always negative, so that the restriction on the auxiliary function can be rewritten *G*_Tb_(*c*) ≤ min {0, −*f′/f*}. Similar constraints on allowable resorption functions *r*_Fm_(*c*) can be developed to ensure that *r*_Tb_(*c*) > 0. Taken together, these conditions give:

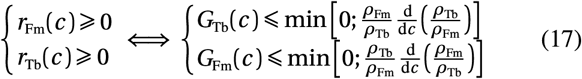

Note that the right-hand sides of these inequalities are only functions of the experimental BMDDs and as such are fully known.

In Figure 5, we build resorption functions *r*_Fm_ and check that the auxiliary function *G*_Fm_ fulfills Eq. (17). In Figure 6, we build an auxiliary function *G*_Tb_ that fulfils Eq. (17), and we determine the corresponding resorption function by Eq. (16). The cortical resorption function *r*_Fm_(*c*) is then found using Eq. (11).

The further away the auxiliary function *G*_Tb_(*c*) is from the upper bound, the further away the minimum resorption rate *r*_Fm_(*c*) is from zero (Ineq. (17)). But also the closer to zero is the corresponding resorption rate *r*_Tb_(*c*) (Eq. (16)). It is clear from this trade-off that the set of physiologically realistic functions *G*_Tb_(*c*) is also bounded from below.

The strict restrictions that exist on the possible choices of the auxiliary functions *G*_Tb_(*c*) imply that the probability to renew interstitial cortical bone would always be very low under Hypothesis 3. This is exemplified in Figure 8a,b, where a region of the qBEI image of one of the femoral cortical samples has been coloured by its corresponding hypothetical tissue age. In this figure, tissue age was calculated from the mineralisation kinetics *c* (*t*) corresponding to the choice of the upper limit of the auxiliary function *G*_Tb_(*c*). Other choices lead to similar tissue age distributions. In all cases, interstitial bone would have to be interpreted as being over 100 years old.

**Figure 8.**
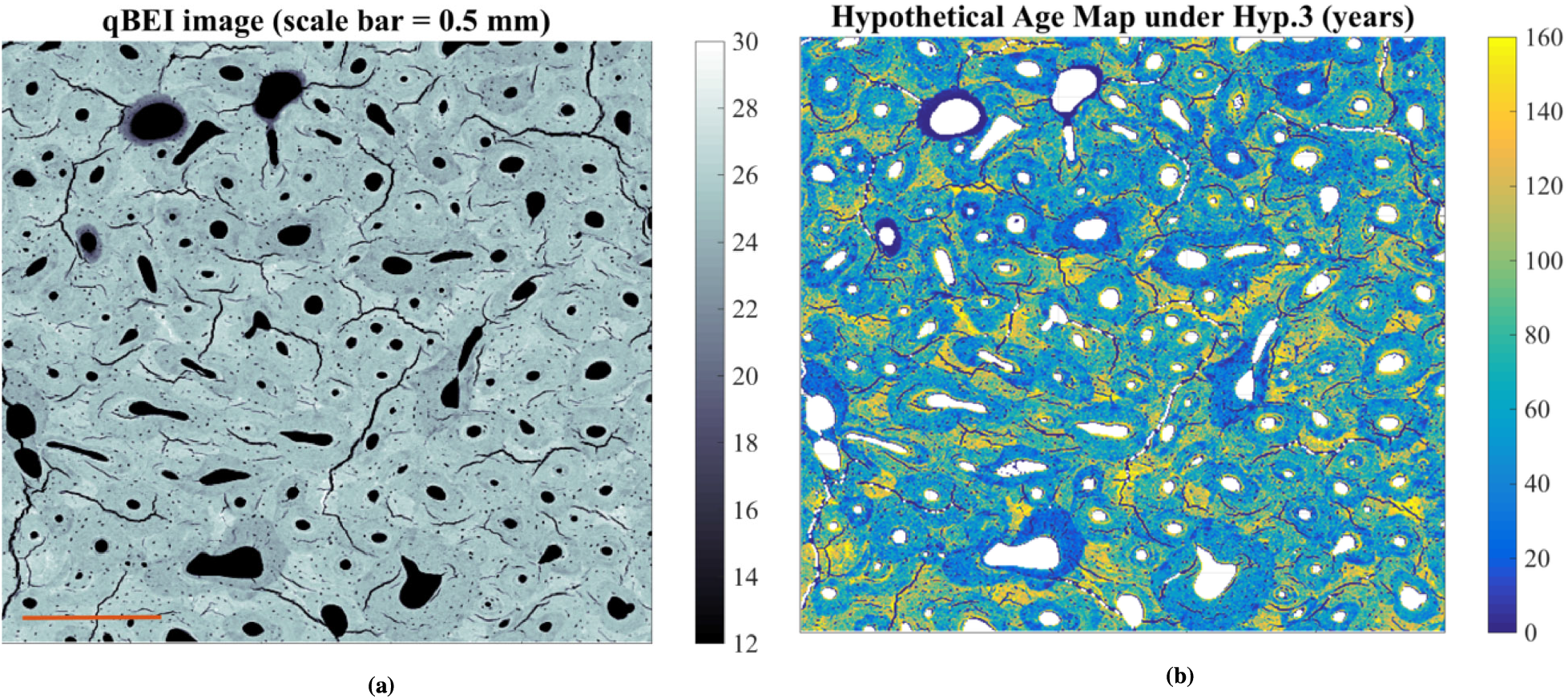
(a) qBEI image, and (b) hypothetical age map resulting from Hypothesis 3, with the cortical resorption function obtained by assuming the auxiliary function *G*_Tb_(*c*) to be the upper bound Condition.

### B.1 Resorption of unmineralised bone

As stated in the Results section, the volume of bone resorbed per unit time comes mainly from the unmineralised bone packets. Here, we give the mathematical steps to reach this statement.

Let *R*(*c\**) be the volume of femoral cortical bone with calcium content less than *c\** resorbed per unit time:

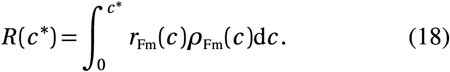

It is clear from Eq. (18) that 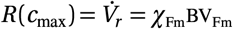. The aim here is to show that under Hypothesis 3, any choice of *r*_Fm_ leads to *B*(*c\**) being unreasonably close to *B*(*c*_max_) already at very small values of *c\**. Let *α* be the fraction of the total volume of bone resorbed per unit time that has a calcium content below *c\**:

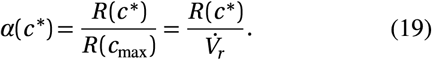

In the following, we provide a lower bound for *α*(*c\**) which we will use to show that under Hypothesis 3 and given the experimental measurements of trabecular and cortical BMDDs, *α*(21 wt% Ca) ≥ 25% and *α*(20 wt% Ca) ≥ 15%. Hypothesis 3 would thus mean that (i) at least 25% of the femoral cortical bone resorbed per unit time would corresponds to bone of calcium content below 21 wt% Ca, which represent the 5% lowest mineralised bone in the femoral cortex; (ii) at least 15% of the femoral cortical bone resorbed per unit time would be resorbed before this bone starts secondary mineralisation, estimated to start at around 20 wt% Ca. These two highly unlikely predictions go against the idea of bone self-repair.

To bound *α*(*c* ^∗^) from below, we first note that 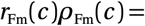 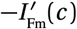 and from Eq. (17),

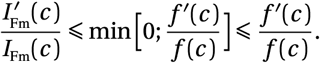

Thus 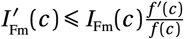, so that

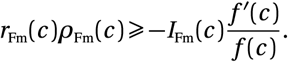

By integrating both sides of the inequality up to *c\**, and using the definition of *R*(*c\**) in Eq. (18), we have:

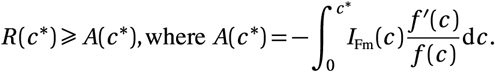

To bound *R*(*c\**) further, we first integrate *A*(*c\**) by parts:

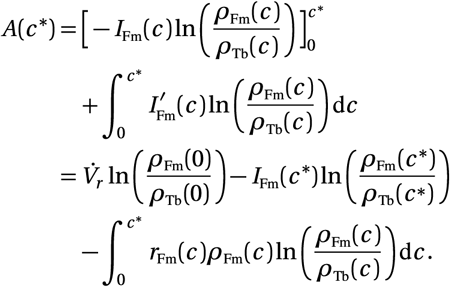

Because the femoral cortical BMDD’s peak is shifted along the calcium axis compared to the trabecular BMDD’s peak, *ρ*_Tb_(*c*) > *ρ*_Fm_(*c*) up until *c* ≃ 24 wt% Ca. Accordingly, the function 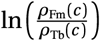 is negative for all *c* ≤ 24 wt% Ca (see Figure 9). Hence, for *c\** < 24wt% Ca,

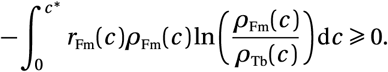

**Figure 9.**
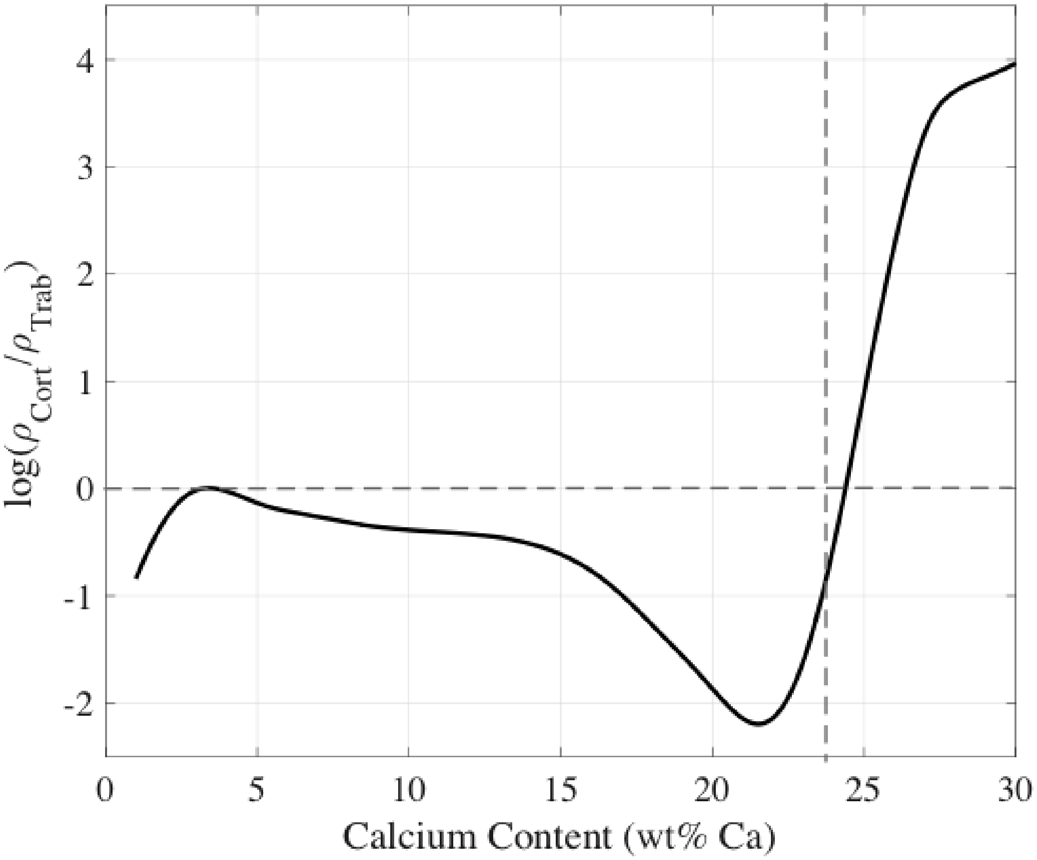
Plot of the function 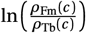, which shows that this function is negative for all *c* ≤ 24 wt% Ca.

This leads to:

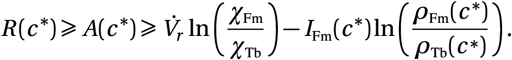

With Eq. (19), this inequality becomes:

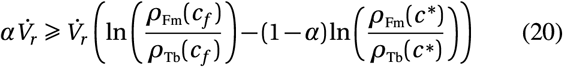

where we used the fact that from the definitions of *I*_Fm_ and *α*, 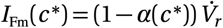. The inequality (20) can be reorganised into

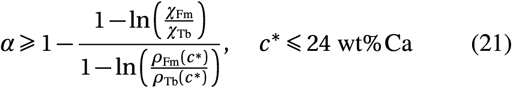

All the quantities in the right hand side of Ineq. (21) are known. Evaluating them at *c\** = 21 wt% Ca and *c\** = 20 wt% Ca shows that *α*(21 wt% Ca) ≥ 25% and *α*(20 wt% Ca) ≥ 15% (Figure 10). In other words, under Hypothesis 3, at least 25% of the total amount of femoral cortical bone resorbed per unit time would have a calcium content less than *c\** = 21 wt% Ca, and at least 15% would have a calcium content less than *c\** = 20 wt% Ca.

**Figure 10.**
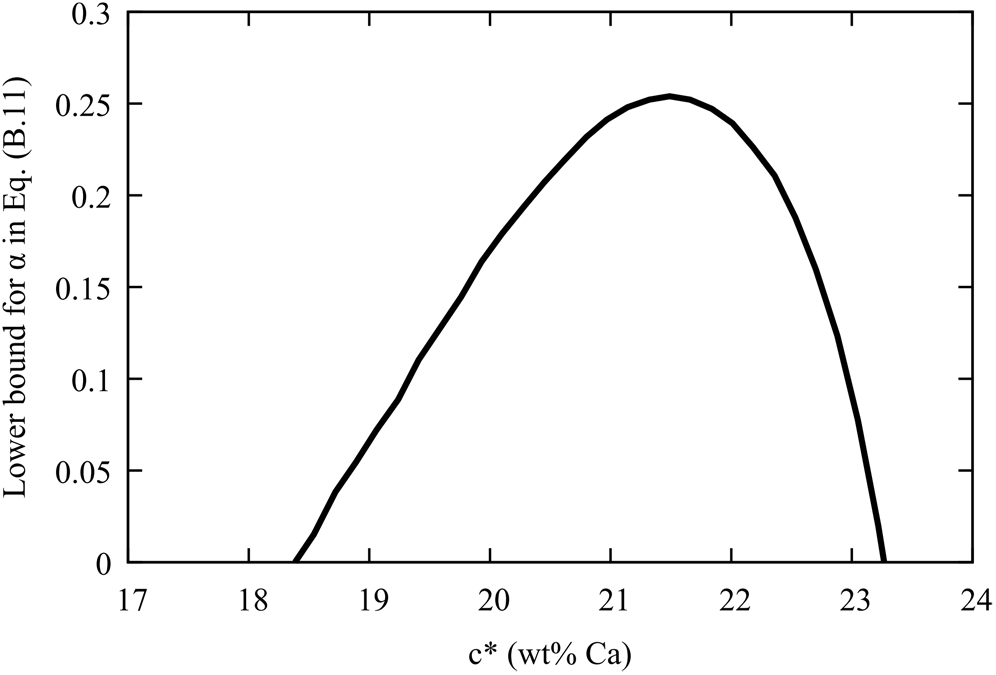
Under Hypothesis 3, the fraction *α*(*c\**) of total volume of bone resorbed per unit time that has a calcium content less than *c\** must be greater than the lower bound in Eq. (21) plotted in this figure. This shows that *α*(21 wt% Ca) ≥ 25% and *α*(20 wt% Ca) ≥ 20%.

## C Goodness of fit of the mineralisation kinetics

To understand the sensitivity of mineralisation kinetic laws to measured BMDDs, we quantify in this appendix how similar two BMDDs have to be in order for their mineralisation kinetics *c*(*t*) to be within the same error as that between the trabecular and the transformed cortical mineralisation kinetics. To weigh the discrepancy between mineralisation kinetic laws by the frequency of occurrence of calcium density, we define the following measure of discrepancy of a mineralisation kinetic law *c*(*t*) compared to that of trabecular bone: 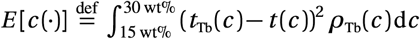, where *t*_Tb_(*c*) and *t*(*c*) are the inverse functions of *c*_Tb_(*t*) and *c*(*t*), respectively. The discrepancy between the scaled cortical mineralisation kinetics and the reference trabecular mineralisation kinetics gives a value *E*_Fm_ = *E*[*c*_Fm_(·)] = 0.017 year^2^. We then vary systematically the reference trabecular BMDD width (Ca_WIDTH_), skewness (Ca_SKEW_), and kurtosis (Ca_KURT_) by applying a parametric transformation following the approach by Jones and Pewsey [34]. For each of these modified trabecular BMDDs, we calculate the corresponding mineralisation kinetics and estimate the error *E* with respect to the reference trabecular mineralisation kinetics. We find that to maintain these errors *E* below *E*_Fm_, Ca_WIDTH_ has to deviate less than 5% from the trabecular value, while skewness and kurtosis may deviate by up to 35%. This shows that the spread of mineral heterogeneity around the mean captured by Ca_WIDTH_ is an important signature of mineralisation kinetics, whereas skewness and kurtosis are less closely related to it.

## D Universal coupling laws between mineralisation kinetics and turnover rate

The very good match between the mineralisation kinetics of trabecular bone and the mineralisation kinetics of femoral cortical bone after a scaling in time and a shift in calcium content in Eqs (1)–(2) suggests that we may define a universal, bone-type-independent mineralisation rate 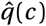 by

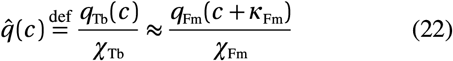

and a universal, bone-type-independent mineralisation kinetic law *ĉ*(*t*) by

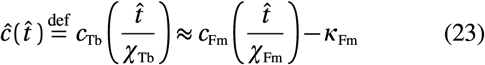

where 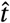 is a dimensionless time variable corresponding to turnover time. The approximations in Eqs (22)–(23) are a direct consequence of the matches observed in Fig. 7, summarised mathematically in Eqs (1)–(2). The functions 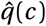 and *ĉ*(*t*) that these matches define in Eqs (22)–(23) are free of bone-type specific time scales, and in that sense, universal. The mineralisation rates and mineralisation kinetic laws in trabecular bone and in femoral cortical bone are retrieved by a simple rescaling in time and shift in mineral density of these scale-free mineralisation laws:

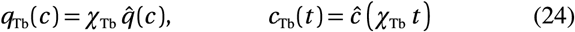

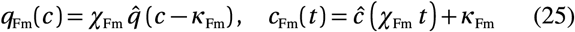

In other words, the time scale of mineralisation in each of these bone types is determined by the time scale of bone turnover only. Equations (24)–(25) summarise the concept that mineralisation is coupled to turnover rate. Equations (22)–(23) are proposed as an explicit experimental test for the validity of this hypothesis. Hypothesis 4 would be further corroborated if measurements of BMDDs in other skeletal sites would fulfil Eqs (22)–(23) for suitable (site-dependent) calcium shifts.

It is important to note here that the scaling laws we observed in Eqs (22)–(23) are completely independent of the values of turnover rate *χ*_Tb_ and *χ*_Fm_ that we have assumed. The division by turnover rate in Eq. (22) cancels the turnover rate dependence of mineralisation rate in Eq. (8). The values assumed in this paper for the trabecular and femoral cortical turnover rate were used in Fig. (7) only to transform the time variable from the dimensionless ‘turnover time’ into absolute units of time (years).

